# Piroplasm parasites (Apicomplexa: Piroplasmida) in northeastern populations of the invasive Asian longhorned tick, *Haemaphysalis longicornis* Neumann (Ixodida: Ixodidae), in the United States

**DOI:** 10.1101/2023.06.26.546593

**Authors:** H Herb, FC Ferreira, J Gonzalez, DM Fonseca

## Abstract

Piroplasms, which include the agents of cattle fever and human and dog babesiosis, are a diverse group of blood parasites of significant veterinary and medical importance. The invasive Asian longhorned tick, *Haemaphysalis longicornis*, is a known vector of piroplasms in its native range in east Asia and invasive range in Australasia. In the US state of Virginia, *H. longicornis* has been associated with *Theileria orientalis* Ikeda outbreaks that caused cattle mortality. We examined 667 *H. longicornis* collected in 2021 from three sites in New Brunswick, New Jersey, the US state where established populations of this species were first detected in 2017. We used primers targeting the 18S small subunit rRNA and the cytochrome b oxidase loci and unveiled the presence of DNA from an unidentified *Theileria* species (in 1 nymph) and *Theileria cervi* type F (1 adult, 5 nymphs). In addition, we sequenced a 130 bp fragment of the cytochrome oxidase b locus from *Odocoileus virginianus*, the white-tailed deer, in a partially engorged questing *H. longicornis*, supporting the association of this tick species with deer. We also detected DNA from an undescribed *Babesia* sensu stricto (‘true’ *Babesia*, 2 adults, 2 nymphs) as well as *Babesia* sp. Coco (1 adult, 1 nymph). Finally, we detected DNA from *Babesia microti* S837 (1 adult, 4 nymphs). *Babesia microti* S837 has been sequenced from striped skunks, *Mephitis mephitis*, and is closely related to the human pathogen *B. microti* US-type. The five parasites we are associating with *H. longicornis* represent a diverse assemblage spanning three clades in the piroplasm phylogeny, two undescribed, raising concerns of transmission amplification of veterinary pathogens as well as spillover of pathogens from wildlife to humans.

## Introduction

Invasive ticks, mosquitoes, and other blood feeding arthropods may introduce and transmit (i.e., vector) exotic pathogens for which local populations have little or no immunity. Resulting disease can range from mild to severe to fatal and can have a significant impact on human health (e.g., Zika fever, West Nile virus encephalitis), animal health (e.g., redwater fever, blue tongue virus), hasten extinction (e.g., bird malaria), and cause economic damage to agriculture, tourism, and other industries (Athni et al., 2021). Invasive vectors can also potentially spread existing wildlife pathogens by creating new transmission pathways, which can have significant ecological and public health implications, particularly in the context of One Health (*sensu* Lerner and Berg, 2015).

The phylum Apicomplexa includes well known blood-borne protozoa such as *Plasmodium falciparum* and *P. vivax*, the primary agents of human malaria (Votýpka et al., 2016). The Apicomplexa class Piroplasmida includes *Babesia, Theileria*, and *Cytauxzoon* that are primarily transmitted by hard ticks (Ixodida: Ixodidae) and can affect a wide range of hosts (Almazán et al., 2022; Onyiche et al., 2021). While piroplasms were once classified based on morphology and host associations alone, the advent of molecular methods has greatly advanced the overall understanding of the diversity of Piroplasmida (Garrett et al., 2019). A recent analysis by Jalovecka et al. (2019) indicates that there are at least 10 distinct clades within Piroplasmida, with both *Babesia* spp. and *Theileria* spp. comprising polyphyletic groups in need of taxonomic revision.

*Babesia* spp. are broadly divided into *Babesia* sensu stricto and *Babesia* sensu lato, with the former representing a monophyletic group considered as “true *Babesia*”, distinguishable from other piroplasms by their ability to infect the reproductive organs of the tick and to be transmitted to the eggs (transovarial transmission) (Jalovecka et al., 2019; Schnittger et al., 2022; Schreeg et al., 2016). Within *Babesia* sensu lato, one of the best-characterized clades is the *Babesia microti* group, which contains the piroplasms responsible for most human babesiosis cases worldwide, especially in the northern United States (Renard and Mamoun, 2021).

Importantly, *Babesia microti* includes at least two different genetic lineages pathogenic to humans and several only known from reservoir hosts such as mice, voles, and skunks (Goethert, 2021). Furthermore, human disease caused by other *Babesia* such as *B. divergens, B. venatorum, B. duncani, B. canis, B. odocoilei*, and *B. bovis* has also been reported (Hong et al., 2019; Scott et al., 2021).

Rates of human babesiosis have been increasing in the US, particularly in the northeastern states (Almazán et al., 2022; Swanson et al., 2023). While those infected with *Babesia* may experience fever, chills, headache, muscle aches, fatigue, and red or brown urine, some may not have any symptoms at all, especially if their immune systems are not compromised (Almazán et al., 2022). People may remain infected for years, and even if asymptomatic, can transmit piroplasms through blood transfusions or organ transplants (Bloch et al., 2019) and transmission from infected mothers to developing fetuses has been demonstrated (Horowitz and Freeman, 2020). Piroplasmid infections are typically treated with various combinations of atovaquone, azithromycin, clindamycin, and quinine, however, concerns regarding side effects, drug resistance, and drug efficacy indicate the need for development of novel treatment options (Renard and Mamoun, 2021).

Bovine babesiosis caused by *Babesia bigemina* and *Babesia bovis* has long been a concern to American cattle ranchers and several *Theileria* species can sicken horses, cervids, and bovids (Almazán et al., 2022; Osbrink et al., 2022). Both babesiosis and theileriosis cause significant economic losses annually to the agricultural industries of many countries due to reduced production, death, abortions, restrictions on animal movement, and costs associated with preventive measures and treatments (Almazán et al., 2022; Dinkel et al., 2021; Osbrink et al., 2022; Schnittger et al., 2022). Companion animals are also at risk, with infections of *Babesia vulpes, Babesia conradae, Babesia vogeli, Babesia gibsoni*, and *Babesia* sp. Coco capable of causing mild to severe disease in dogs in the United States (Dear and Birkenheuer, 2022).

Since the initial discovery of the invasive Asian longhorned tick (*Haemaphysalis longicornis*) in the United States in 2017 (Rainey et al., 2018), there have been concerns regarding the potential threats this ectoparasite may pose. In North America, as in Australasia where it expanded to in the early 20^th^ century, *H. longicornis* reproduces asexually (clonally) by parthenogenesis (Schappach et al. 2020), which underlies the ability of this species to develop large populations very quickly. In its Australasian range, *H. longicornis* represents a major threat to domestic livestock, heavily parasitizing large ruminants, and impeding production (Heath, 2016).

Globally, *H. longicornis* is a known vector of piroplasms that infect humans, livestock, and companion animals, including *B. ovata, B. gibsoni, B. microti, T. uilenbergi*, and *T. orientalis* (Dear and Birkenheuer, 2022; Dinkel et al., 2021; Gray et al., 2019; Li et al., 2009; Wu et al., 2017) and may also be a vector of *Babesia caballi*, the agent of equine babesiosis (Bautista et al., 2001). As in Australia (Marendy et al., 2020), in Virginia, USA, *H. longicornis* is implicated in the transmission of the virulent *Theileria orientalis* Ikeda genotype that resulted in multiple cattle deaths (Dinkel et al., 2021; Thompson et al., 2020).

Although humans are not favored hosts of *H. longicornis*, opportunistic feeding is well documented both in the native and invasive ranges (Bickerton and Toledo, 2020; Wormser et al., 2020). In East Asia, *H. longicornis* vectors severe fever with thrombocytopenia syndrome virus (SFTSV), an emerging human tick-borne disease recently reclassified as Dabie bandavirus (Li et al., 2021; Liu et al., 2015; Luo et al., 2015). Under laboratory conditions, US lineages of *H. longicornis* can vector the closely related Heartland virus (Raney et al., 2022b) as well as Powassan virus (Raney et al., 2022a), two native pathogenic viruses emergent in parts of the US. They can also vector *Rickettsia rickettsii*, the causative agent of Rocky Mountain Spotted Fever (Stanley et al., 2020). In addition, Bourbon virus has been detected from a larval pool, two nymphs and one adult field collected *H. longicornis* in Virginia, USA (Cumbie et al., 2022).

While *H. longicornis* is currently not perceived as a major public health threat in the US, this status may change given the enormous densities it can reach in favorable habitats (Bickerton et al., 2021; González et al., 2023; Rochlin et al., 2023; Schappach et al., 2020).

Our objective was to assess the potential role of *H. longicornis* as a vector of piroplasms in NJ, the most urbanized US state that, maybe surprisingly to many, also boasts the highest density of horses (Rankins and Malinowski, 2020).

## Methods

### Study Areas

This study was conducted in three sites approximately 1.2 km from each other within the Rutgers University Cook Campus in New Brunswick, NJ (please refer to Ferreira et al., in press, for a map). Surveys for *H. longicornis* were initiated at these sites in 2018 when the species was first detected on a grassy area next to a goat pen (Egizi et al., 2019a), a site that became known as the “Goat Farm” (40.47444° N, 74.43683° W). The “Rutgers Gardens” site (40.47455° N, 74.42030° W) is inside a 180-acre botanical garden, consisting of designed gardens, plant collections and natural habitats. Finally, the “University Inn” site (40.48413 N, 74.43051 W) is a meadow and forested park behind the Rutgers University Inn & Conference Center. At all sites, local forest is dominated by oak and maple trees and huckleberry and blueberry shrubs (Breden et al., 2001), with grassy ecotones.

### Tick surveillance

From June through September 2021, concomitant with surveys for ticks on mammals at the same sites, we collected questing ticks flagging between 50-75 m^2^ per sampling session at each site (Ferreira et al., in press). Tick sampling was performed using a white crib flannel sweep measuring 50 x 100 cm with a PVC pipe handle (Egizi et al., 2019a). The sweep cloth was checked in 1-2 m intervals since *H. longicornis* does not attach firmly to the flannel and often drops off over longer intervals (Bickerton et al., 2021). Sampling was conducted by sweeping over the vegetation. Ticks were collected from both sides of the flag and morphologically identified in the laboratory to the species level using a stereomicroscope (Leica S8 APO, Leica Microsystems) following appropriate taxonomical keys (Egizi et al., 2019b; Keirans and Litwak, 1989). The larvae of *H. longicornis* were not stored during these surveys and were not available for pathogen testing. A few questing *H. longicornis* that were found partially engorged (sensu Price et al., 2022) were processed separately (see section below on “Bloodmeal analysis of partially engorged specimens”).

### DNA extraction and pathogen detection

We placed each tick in 180 µl of Qiagen buffer ATL with 20 µl Qiagen Proteinase K (10mg/ml) in microfuge tubes and homogenized them with 5 mm sterile glass beads (Fisher Scientific, Waltham, MA) using a TissueLyser bead mill (Qiagen Inc., Valencia, CA, USA). We extracted DNA from individual ticks using Dneasy Blood and Tissue single column kits (Qiagen Inc., Valencia, CA, USA) following the manufacturer’s instructions. We eluted DNA from each column twice with 50 µl of Qiagen’s elution buffer AE into separate labelled microtubes.

After reviewing the literature, we chose primers targeting the multi copy 18S rRNA locus (**Table 1**) to match a broad range of piroplasm species (Casati et al. 2006) and tested all ticks individually. To further characterize a putatively new *Babesia* sp., we tested an infected specimen of *H. longicornis* using primers targeting the mitochondrial cytochrome oxidase b (*cytb*) locus (**Table 1**) shown to work across multiple *Babesia* species (Wickramasekara Rajapakshage et al. 2012).

**Table 1.**
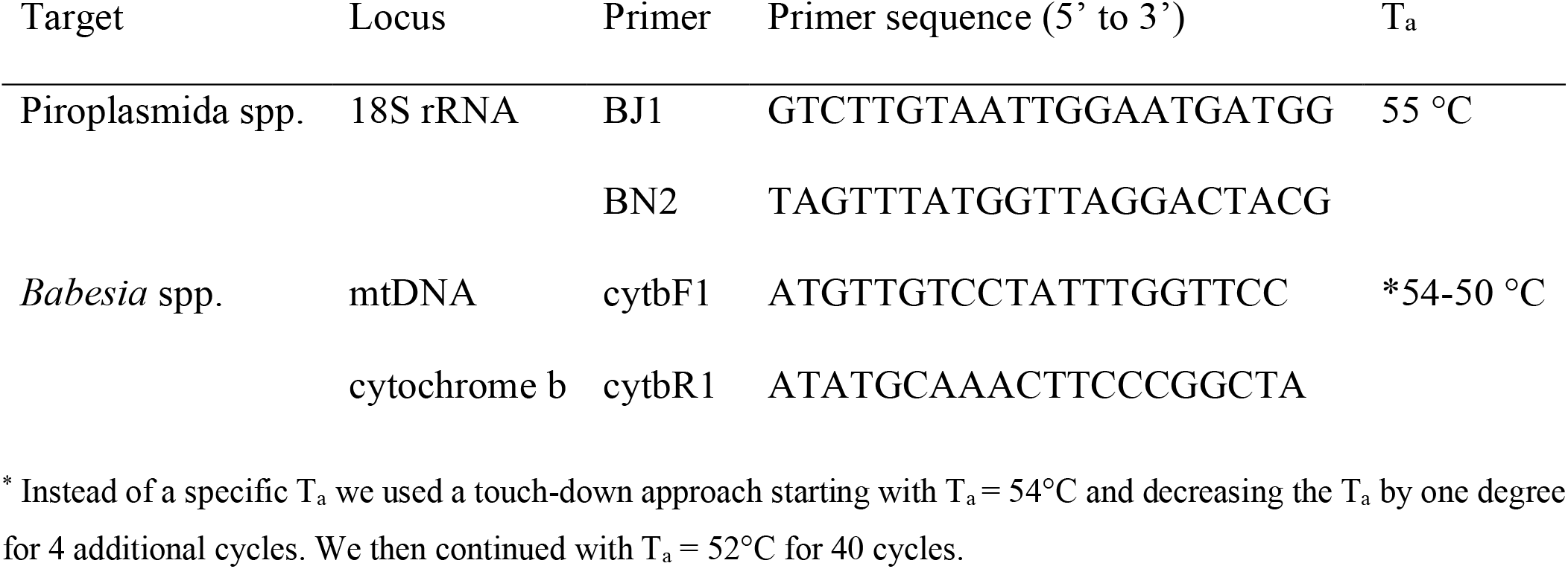
Primers used for piroplasm detection in *H. longicornis*. T_a_ = annealing temperature.

We amplified the targeted loci in 20 µl reactions with Amplitaq Gold Master Mix (ThermoFisher Scientific, Waltham, MA, USA) following the manufacturer’s protocol. After visualizing the amplification in a 1% agarose gel, we cleaned the PCR reactions with ExoSAP-IT (ThermoFisher Scientific, Waltham, MA, USA) and performed Sanger sequencing separately with both primers at Azenta Genewiz (South Plainfield, NJ, USA). The sequences were trimmed and aligned with Geneious Prime 2023.0.1 (Biomatters Inc. San Diego, CA, USA) and the consensus was used as a query in NCBI’s Basic Local Alignment Search Tool, BLASTn (Altschup et al., 1990).

### Phylogenetic Analysis

Consensus sequences were aligned to available NCBI Genbank sequences (Benson et al. 2012) representative of the major piroplasm clades (Jalovecka et al., 2019) and the alignments were trimmed to the same size (557 bp). We constructed maximum likelihood phylogenetic trees based on the 18S rRNA and *cytb* loci using IQ-TREE with 1,000 ultrafast bootstrap replicates (Hoang et al., 2018; Nguyen et al., 2015). ModelFinder was used to choose the best-fitting substitution model based on Bayesian Information Criterion (Kalyaanamoorthy et al., 2017).

*Cardiosporidium cionae* (GenBank acc. Num. EU052685) and *Babesia microti* (GenBank acc. Num. NC034637) were used as outgroups for the 18S rRNA and *ctyb* phylogenetic trees, respectively.

### Statistical Analysis

To determine whether there statistically significant differences in infection rates in nymphal vs. female *H. longicornis*, we used a binomial generalized linear model using the glm() function in R (R Core Team, 2022). To determine the correlation between sample size and infection rates, we used the lm() function in R (R Core Team, 2022).

### Bloodmeal analysis of partially engorged specimens

We isolated DNA from two partially engorged *H. longicornis* collected on July 15, 2021, (one nymph and one adult) using DNeasy Blood and Tissue columns (Qiagen, Valencia, CA).

We included an extraction control, and all work was performed in a dedicated clean lab inside a laminar flow hood (Mystaire, Creedmor, NC, USA). We used primers CytbVertR1 (Egizi et al., 2013) and BMF1 (originally called H15149, Kocher et al., 1989) to amplify a 132 nucleotide fragment in the cytochrome oxidase b locus. A PCR product obtained from the adult *H. longicornis* was purified (ExoSAP-IT, Affymetrix, Santa Clara, CA) then sequenced at Azenta Genewiz (South Plainfield, NJ, USA). The sequences were trimmed and aligned with Geneious Prime 2023.0.1 (Biomatters Inc. San Diego, CA, USA) and the consensus was used as a query in NCBI’s Basic Local Alignment Search Tool, BLASTn (Altschup et al., 1990).

### In silico *tests of existing qPCR assays against* B. microti S837

We identified six published real-time PCR (rtPCR) assays specific for *B. microti* targeting the 18S region (Bloch et al., 2013; Hersh et al., 2012; Hojgaard et al., 2014; Rollend et al., 2013; Teal et al., 2012; Tonnetti et al., 2009). We then performed an *in silico* analysis by comparing the primer and probe sequences from the publications to an alignment of (1) a 18S sequence of *B. microti* S837 from GenBank (AY144698, 1255 bp), (2) the sequences we recovered from *H. longicornis* (457 bp) and (3) two sequences in Genbank that are considered representative of *B. microti* US-type (AB085191, 1721 bp and AB190459, 2536 bp). The oligonucleotides were mapped to the target sequences using Geneious Prime 2023.0.1 (Biomatters Inc. San Diego, CA, USA).

## Results

We screened 667 *H. longicornis* nymphs and adults collected from the environment and found evidence of piroplasm parasites in 18 ticks (2.7%, **Table 2**). Adult and nymph infection rates did not differ (infection rates of 2.4% and 2.8%, respectively, p-value = 0.60). Infection rates among sampling sites reflected sample size (r^2^ = 0.92, p-value <0.01; using adult and nymph data separately to increase the statistical power).

**Table 2.** Number of *H. longicornis* nymphs and adults collected from the environment at three sites at Rutgers Cook campus and tested for piroplasm parasites (# Pos represents the number of ticks that were positive for piroplasm DNA). Collections were started on 24 June 2021 and proceeded approximately bi-weekly until 10 September 2021 (refer to Ferreira et al., in press, for details).

|  | June |  | July |  | August |  | September |  | Total |  |
| --- | --- | --- | --- | --- | --- | --- | --- | --- | --- | --- |
|  | N | # Pos | N | # Pos | N | # Pos | N | # Pos | N | # Pos |
| **Goat Farm** | **35** | **2** | **100** | **4** | **14** | **0** | **4** | **0** | **153** | **6** |
| Nymph | 20 | 2<sup>a</sup> | 38 | 0 | 7 | 0 | 1 | 0 | 66 | 2 |
| Adult | 15 | 0 | 62 | 4<sup>abc</sup> | 7 | 0 | 3 | 0 | 87 | 4 |
| **Rutgers Gardens** | **26** | **0** | **30** | **0** | **8** | **0** | **0** | **0** | **64** | **0** |
| Nymph | 22 | 0 | 16 | 0 | 4 | 0 | 0 | 0 | 42 | 0 |
| Adult | 4 | 0 | 14 | 0 | 4 | 0 | 0 | 0 | 22 | 0 |
| **University Inn** | **78** | **1** | **308** | **10** | **56** | **1** | **8** | **0** | **450** | **12** |
| Nymph | 64 | 1<sup>c</sup> | 250 | 9<sup>abcde</sup> | 33 | 1<sup>a</sup> | 7 | 0 | 354 | 11 |
| Adult | 14 | 0 | 58 | 1<sup>d</sup> | 23 | 0 | 1 | 0 | 96 | 1 |
| **Grand Total** | **139** | **3** | **438** | **14** | **78** | **1** | **12** | **0** | **667** | **18** |

<sup>d</sup> *Theileria cervi* (a total of 6 positive ticks)
<sup>e</sup> *Theileria* sp. (1 positive ticks)
<sup>b</sup> *Babesia* sp. Coco (2 positive ticks)
<sup>c</sup> *Babesia* sp. (4 positive ticks)
<sup>a</sup> *Babesia microti* S837 (5 positive ticks)

We did not see evidence of piroplasm coinfections (such as double chromatogram peaks) in the positive ticks. The primers targeting the 18S rRNA gene amplified fragments ranging in size from 395 to 515 base pairs (bp) spanning the V4 hypervariable region (Cauvin et al., 2019).

The 18S rRNA fragment from a nymph collected at University Inn had a 100% pairwise identity to five *Theileria* sp. sequences (GenBank acc. nums. MW008536, MW008531, MK262962, MK262963, and MK262959) that are considered “Type X” or “divergent” (Cauvin et al., 2019; Olafson et al., 2020). This sequence differed by 20 bp (17 mismatches and 3 deletions) from six other sequences obtained from *H. longicornis* also collected from the University Inn site. The closest match for these six sequences (99.8% to 100% pairwise identity) was a *Theileria cervi* type F sequence (GenBank U97054). Three of the six sequences, two from nymphs and one from an adult, were identical to U97054 while the remaining three, all from nymphs, differed by one bp (GenBank acc. num **TBD**). All sequences clustered within a clade containing *Theileria cervi* and *Theileria orientalis* (**Figure 1a**).

**Figure 1.**
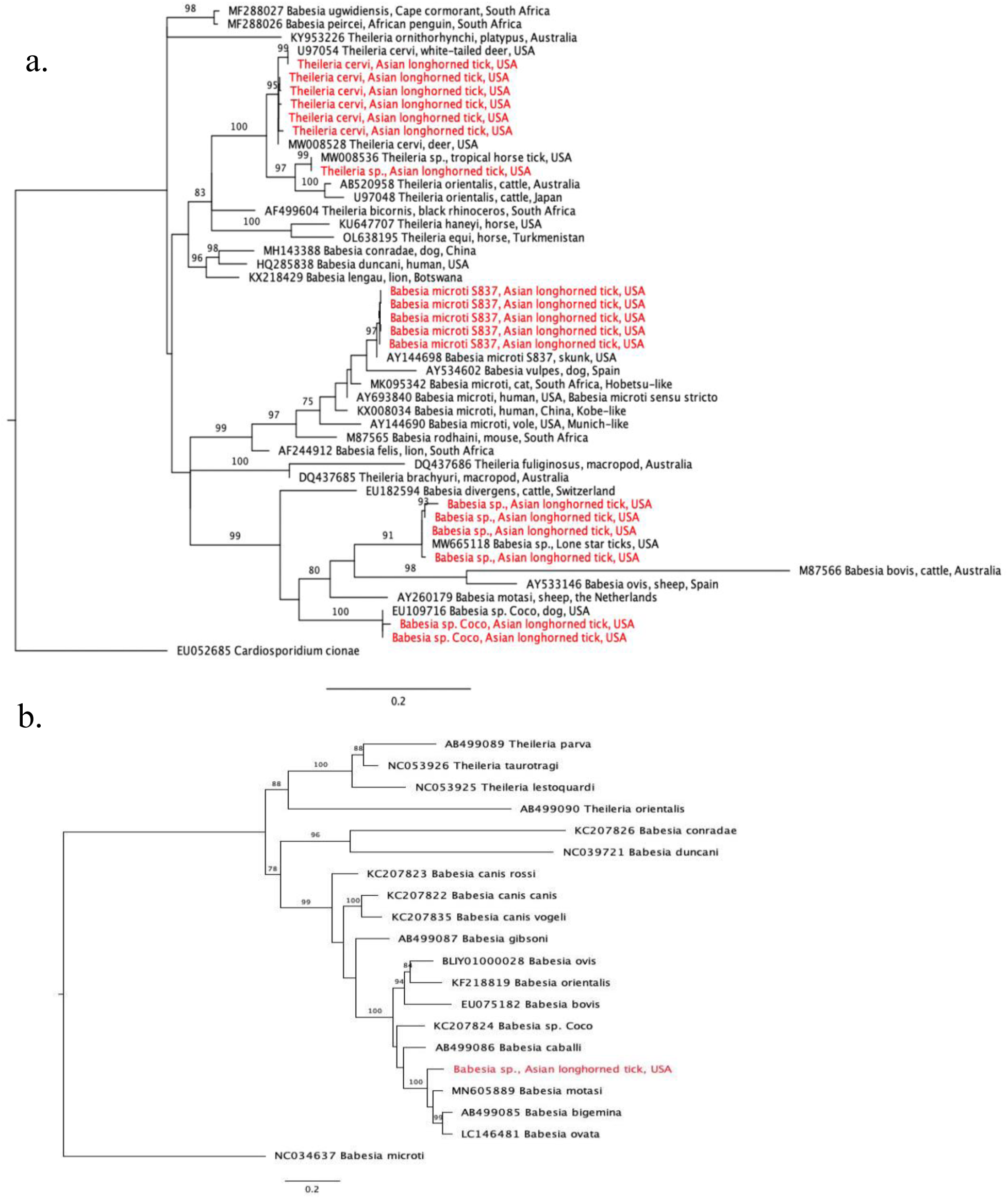
Phylogenetic trees with piroplasm sequences obtained from *Haemaphysalis longicornis* (in red). a.) Tree based on 18S rRNA locus. Constructed with TIM2+F+I+G4 substitution model. b.) Tree based on cytochrome b locus. Constructed with K3Pu+F+I+G4 substitution model.

Two 18S rRNA sequences, one from a Goat Farm adult and the other from a University Inn nymph, were identical to a *Babesia* sp. isolate NYT-435 (GenBank acc. num. MW665118) recently sequenced from a pool of lone star ticks (*Ambylomma americanum*) from Staten Island, NYC (Jain et al., 2021). The sequence of the *Babesia* sp. isolate NYT-435 was also the closest match for two other sequences, one from another Goat Farm adult and a second University Inn nymph (99.7% and 99.5% pairwise identities with a one bp and two bp difference, respectively; GenBank acc. num **TBD**). The next best match for these four sequences (97.9% to 98.2% pairwise identity) was a *Babesia* sp. from a white-tailed deer in Texas (GenBank acc. num.

HQ264120). These four sequences clustered within the *Babesia* sensu stricto clade with *Babesia bovis* and *Babesia ovis* (**Figure 1a**). From one of the specimens that were positive for the unknown *Babesia*, we also amplified a 1,009 bp DNA fragment with the *cytb* primers (GenBank acc. num **TBD**). The closest match to this sequence was *Babesia motasi* isolate Lintan (GenBank acc. num. MN605889) with 90.8% pairwise identity. The sequence from this *H. longicornis* clustered with *Babesia motasi* in the *Babesia* sensu stricto clade (**Figure 1b**).

The 18S rRNA from two sequences, one from a University Inn nymph and another from a Goat Farm adult, matched *Babesia* sp. Coco. The sequence from the Goat Farm adult was 100% pairwise identical to Genbank EU109716, whereas the sequence from the University Inn nymph differed by one bp and had a 99.8% pairwise identity (GenBank acc. num **TBD**).

Finally, five 18S rRNA sequences, from two nymphs and one adult from the Goat Farm and two nymphs from the University Inn, had a 100% pairwise identity to *Babesia microti* isolate S837 (GenBank acc. num. AY144698). In our analysis, these sequences clustered with *Babesia vulpes* within the *Babesia microti*-like group (**Figure 1a**, Jalovecka et al. 2019).

The *in silico* examination of the six rtPCRs developed to identify *Babesia microti* US-type revealed that two were a 100% match to *B. microti* S837 (Hersh et al., 2012, Hojgaard et al., 2014). The reverse primer (Bm18Sr) and the probe (Bm18Sp) of a third assay developed by Rollend et al. (2013) were also a 100% match to *B. microti* S837. The forward primer (Bm18Sf) showed a single nucleotide difference towards the 5’ end, which is unlikely to diminish its effectiveness as a primer for *B. microti* S837. Unfortunately, the assays developed by Bloch et al. (2013), Tonnetti et al. (2009) and Teal et al. (2012) target regions of the 18S locus outside the available sequences for *B. microti* S837.

We amplified and sequenced a 132 bp DNA fragment from a partially engorged adult *H. longicornis* that matched the cytb locus of white-tailed deer, *Odocoileus virginianus* (GenBank Accession# AF535863).

## Discussion

We report the detection of DNA sequences belonging to three piroplasm clades in 2.7% field collected *Haemaphysalis longicornis* nymphs and adults in New Jersey, United States.

Infection rates for *Theileria, Babesia* sensu stricto, and *Babesia microti*, were 1%, 0.9% and 0.8%, respectively. While these may be low piroplasm infection rates compared to studies that detected *Theileria orientalis* in 12.7% of *H. longicornis* in Virginia (Thompson et al. 2020, 2021), those were collected from the cattle farm where the first US outbreak of *T. orientalis* Ikeda occurred, which would have increased the likelihood that local vectors were infected.

Overall, there have been few exploratory examinations of piroplasms in *H. longicornis* in the United States.

This is the first report of *Theileria* species besides *Theileria orientalis* Ikeda in *H. longicornis* in the United States. We detected sequences matching *Theileria cervi* Type F, a species that commonly infects deer and other cervids (Cauvin et al. 2019, Olafson et al. 2020) and a sequence that falls within the *Theileria* clade but differs in at least 20 bp from *T. cervi* Type F and matches a strain denoted “divergent” or “type X” (Olafson et al., 2020). The “divergent” or “Type X” strain has been found in wild and farmed deer in Florida and in *Anocenter nitens*, the tropical horse tick, parasitizing white-tailed deer in Texas (Cauvin et al., 2019; Olafson et al., 2020). US populations of *H. longicornis* have often been reported feeding on white-tailed deer (Tufts et al. 2020, USDA, 2023), and our finding of deer DNA in a partially engorged tick supports this. We conclude that *H. longicornis* may be involved in the transmission cycle of *Theileria* in NJ.

To our knowledge, this is also the first report of *Babesia* in questing un-engorged *H. longicornis* in the United States. Specifically, the phylogenetic analyses using both 18S and cytochrome b loci indicate the unknown *Babesia* sequences fall within the *Babesia* sensu stricto clade. *Babesia* sensu stricto (also referred to as “true *Babesia*”) are pathogens of both veterinary and medical importance capable of transovarial transmission in their tick vectors, allowing the parasite to propagate in the absence of vertebrate reservoirs (Jalovecka et al. 2019).

Furthermore, we detected *Babesia* sp. Coco, another “true *Babesia*” in two *H. longicornis. Babesia* sp. Coco can be pathogenic to dogs but usually only if they are immunocompromised (Birkenheuer et al., 2004; Dear and Birkenheuer, 2022; Holman et al., 2009; Sikorski et al., 2010), so it is unclear whether dogs are incidental hosts or are an important reservoir species that become symptomatic when immunocompromised. *Haemaphysalis longicornis* have been reported feeding on dogs in the United States (Thompson et al., 2022; Trout Fryxell et al., 2021).

Finally, to the best of our knowledge, this is the first report of *Babesia microti* genotype S837 in *H. longicornis*. This genotype is found in skunks *Mephitis mephitis* (Goethert, 2021). In our phylogenetic analysis, *B. microti* S837, our sequences, and *Babesia vulpes* form a sister group to members of the other *B. microti* clades **(Figure 1**). Despite clustering outside of the clade containing US-type *B. microti, B. microti* S837 *in silico* analysis revealed that three qPCR assays commonly used to detect *B. microti* US-type in both human samples and ticks (Hersh et al., 2012, Hojgaard et al., 2014, and likely Rollend et al., 2013) will amplify *B. microti* S837.

These results indicate that the existing diversity of *Babesia* infecting humans in the US may be underestimated by current diagnostic practices (Goethert, 2021). Indeed, there is an overall lack of knowledge of the biology and epidemiology of wildlife piroplasms in the northeastern US. In areas endemic for human babesiosis molecular studies of wildlife piroplasms rarely employ sequencing to confirm parasite identity, although piroplasm parasites thought to only infect wildlife and/or domestic animals have recently been reported infecting humans in North America (Scott et al., 2021) and elsewhere (Hong et al. 2019). Furthermore, *H. longicornis* has been shown to be a competent vector of *B. microti* under experimental conditions (Wu et al., 2017) and skunks are an important host for this tick species in NJ (Ferreira et al., in press).

Although the health risk of *H. longicornis* to US livestock was established by the outbreak of *T. orientalis* Ikeda in cattle in the state of Virginia (Dinkel et al., 2021; Thompson et al., 2020), there is limited research regarding the veterinary and medical significance of *H. longicornis* in the US, which has focused primarily on testing the ability of US specimens to transmit pathogens of known public health concern (Breuner et al., 2020; Raney et al., 2022b, 2022a; Stanley et al., 2020). By discovering various piroplasm parasites in questing *H. longicornis* specimens from New Jersey, our investigation brings forth new concerns. As a result, it becomes crucial to prioritize studies that delve into the role of *H. longicornis* as an actual vector for these pathogens.

## Acknowledgments

We thank Nicole Wagner MS from the Rutgers Center for Vector Biology and Julia Brennan, now developing her PhD at Virginia Tech, for their technical support. We also thank Dr. Glen Scoles, USDA-ARS Beltsville, for sharing his piroplasm expertise.

## Data availability Statement

Nucleotide sequence data reported in this paper will be made available in the GenBank™, EMBL and DDBJ databases under the accession numbers **TOBEDETERMINED**

## CRediT authorship contribution statement

**Heidi E. Herb:** Conceptualization, Methodology, Formal analysis, Investigation, Data curation, Writing – original draft, Writing - Review & Editing, Visualization. **Francisco C. Ferreira:** Conceptualization, Investigation, Data curation, Writing - Review & Editing. **Julia González:** Conceptualization, Investigation, Data curation, Writing - Review & Editing. **Dina M. Fonseca:** Conceptualization, Methodology, Resources, Writing - Review & Editing, Supervision, Project Administration, Funding Acquisition.

